# Oxygenation properties of hemoglobin and the evolutionary origins of isoform multiplicity in an amphibious air-breathing fish, the blue-spotted mudskipper (*Boleophthalmus pectinirostris*)

**DOI:** 10.1101/818807

**Authors:** Jay F. Storz, Chandrasekhar Natarajan, Magnus K. Grouleff, Michael Vandewege, Federico G. Hoffmann, Xinxin You, Byrappa Venkatesh, Angela Fago

**Author notes:** Department of Biology, Eastern New Mexico University, Portales, NM 88130, USA.

## Abstract

Among the numerous lineages of teleost fish that have independently transitioned from obligate water-breathing to facultative air-breathing, evolved properties of hemoglobin (Hb)-O_2_ transport may have been shaped by the prevalence and severity of aquatic hypoxia (which influences the extent to which fish are compelled to switch to aerial respiration) as well as the anatomical design of air-breathing structures and the cardiovascular system. Here we examine the structure and function of Hbs in an amphibious, facultative air-breathing fish, the blue-spotted mudskipper (*Boleophthalmus pectinirostris*). We also characterized the genomic organization of the globin gene clusters of the species and we integrated phylogenetic and comparative genomic analyses to unravel the duplicative history of the genes that encode the subunits of structurally distinct mudskipper Hb isoforms (isoHbs). The *B. pectinirostris* isoHbs exhibit high intrinsic O_2_-affinities, similar to those of hypoxia-tolerant, water-breathing teleosts, and remarkably large Bohr effects. Genomic analysis of conserved synteny revealed that the genes that encode the α-type subunits of the two main adult isoHbs are members of paralogous gene clusters that represent products of the teleost-specific whole-genome duplication. Experiments revealed no appreciable difference in the oxygenation properties of co-expressed isoHbs in spite of extensive amino acid divergence between the alternative α-chain subunit isoforms. It therefore appears that the ability to switch between aquatic and aerial respiration does not necessarily require a division of labor between functionally distinct isoHbs with specialized oxygenation properties.

**Summary statement:** The blue-spotted mudskipper routinely switches between aquatic and aerial respiration. This respiratory versatility is associated with properties of hemoglobin-oxygen transport that are similar to those found in hypoxia-adapted water-breathing fishes.

## INTRODUCTION

Facultative air-breathing fishes have always been of interest to comparative physiologists because of the versatility required to switch between aquatic and aerial respiration (Graham, 1997; Graham and Wegner, 2010; Bayley et al., 2019). Moreover, the physiological features that distinguish air-breathing fishes from their obligate water-breathing kin may provide clues to the types of phenotypic changes that facilitated the invasion of land by the shallow-water progenitors of modern tetrapods. In addition to evolutionary changes in branchial and cardiovascular function in fish that have evolved the capacity for aerial respiration, there has been much interest in associated changes in hemoglobin (Hb) function and respiratory gas transport (Damsgaard et al., 2014; Damsgaard et al., 2015; Bayley et al., 2019).

Regardless of breathing mode, the physiologically optimal Hb-O_2_ affinity is dictated by the trade-off between O_2_-loading at the respiratory surfaces (Sgills, lungs, or other air-breathing structures) and O_2_ unloading to respiring tissues (Brauner and Wang, 1997; Wang and Malte, 2011), and there are reasons to expect that this trade-off may be especially profound in facultative air-breathing fishes. Since air has a much higher O_2_ content than water, aerial breathing should generally place a lower premium on O_2_ uptake at the respiratory surfaces (since arterial O_2_ saturation is typically not a limiting factor in tissue O_2_ delivery). This has led some authors to suggest that the evolutionary transition from water- to air-breathing generally entailed a reduction in Hb-O_2_ affinity that permitted a concomitant increase in circulatory O_2_ delivery and aerobic metabolism (Johansen and Lenfant, 1972; Johansen et al., 1978; Powers et al., 1979). However, among amphibious fish that alternate between breathing modes, a high Hb-O_2_ affinity retains much utility under conditions of aquatic hypoxia because it improves O_2_ uptake at the gills, thereby enhancing blood O_2_ capacitance (the quantity of O_2_ unloaded to the tissues for a given difference in arterial and venous partial pressures of O_2_ [*P*O_2_]). Indeed, among water-breathing and facultative air-breathing fish alike, increased Hb-O_2_ affinities are generally associated with adaptation to aquatic hypoxia (Powers, 1980; Jensen, 2004; Mandic et al., 2009; Wells, 2009; Fago, 2017; Harter and Brauner, 2017).

In addition to considerations related to differences in O_2_ capacitance between air and water and the challenges posed by aquatic hypoxia, the optimal Hb-O_2_ affinity in air-breathing fish may also relate to the diffusive conductance of air-breathing structures and design features of the cardiovascular system. Among most air-breathing teleosts, extrabranchial gas exchange occurs across the vascularized epithelia of the buccopharyngeal cavity, esophagus, intestine, stomach, or swim bladder (Graham, 1997; Graham and Wegner, 2010). Such air-breathing structures generally have much lower diffusion capacities and lower gas exchange efficiencies than normal gills, so an elevated Hb-O_2_ affinity can promote O_2_ uptake by maintaining a steep *P*O_2_ gradient across the respiratory epithelium (Hlastala and Berger, 2001; Damsgaard et al., 2014). Finally, aerial breathing generally increases the *P*O_2_ of arterial blood in the ventral aorta that flows towards the gills and, in the absence of a non-respiratory branchial shunt, O_2_-rich arterial blood could desaturate during branchial passage (Randall et al., 1981; Olson, 1994; Ishimatsu, 2012; Bayley et al., 2019). Thus, a high Hb-O_2_ affinity may also be beneficial in facultative air-breathers because it helps prevent transbranchial O_2_ loss, thereby ensuring adequate O_2_ delivery to metabolizing tissues. For this reason, Damsgaard et al. (2014) suggested that facultative air-breathing fish species that do not possess transbranchial shunts could be expected to have evolved higher Hb-O_2_ affinities than obligate air-breathers with reduced gills.

Most teleost fish co-express multiple, structurally distinct isoHbs in adult red blood cells, which are conventionally classified as “anodic” or “cathodic” on the basis of electrophoretic mobility (Weber, 1990; Weber, 2000). Anodic isoHbs tend to have relatively low O_2_-affinities and a large Bohr effect (reduced Hb-O_2_ affinity at low pH), whereas cathodic isoHbs tend to have relatively high O_2_-affinities, a heightened sensitivity to the affinity-reducing effects of organic phosphates (e.g., ATP and GTP), and a negligible Bohr effect in the presence of organic phosphates (Weber and Jensen, 1988; Weber, 1990; Jensen et al., 1998; Weber, 2000; Weber et al., 2000; Wells, 2009). According to the classification scheme of Weber (1990), “class I” species like plaice and carp express multiple anodal isoHbs with very similar O_2_-binding properties, whereas “class II” species like anguillid eels, salmonids, and catfish additionally express one or more cathodal isoHbs that broaden the spectrum of O_2_-affinities and allosteric regulatory capacities. The isoHb multiplicity in class II species may permit a physiological division of labor whereby pH-insensitive cathodal isoHbs provide a reserve capacity for blood-O_2_ transport under conditions of environmental hypoxia or metabolic acidosis (Weber, 1990; Weber, 2000). In some teleost species, evidence suggests that regulatory switches in isoHb expression may play a role in the acclimatization response to environmental hypoxia (Rutjes et al., 2007) and in amphibious fish like mudskippers it seems plausible that a physiological division of labor between isoHbs with different oxygenation properties could contribute to the versatility required to reversibly switch between aquatic and aerial respiration.

Here we examine the structure and function of Hbs in the blue-spotted mudskipper (*Boleophthalmus pectinirostris*), an amphibious, facultative air-breathing fish that is distributed along the coastlines of China, Taiwan, the Korean peninsula, and Japan in the northwest Pacific. This species routinely switches between breathing modes when shuttling between tidepools and mudflats in the intertidal zone (Martin and Bridges, 1999) and it uses an extensive respiratory surface for aerial breathing, including the buccal, pharyngeal, branchial, and opercular cavities in addition to the gills and integument (Graham, 1997). We also report sequence data for the full complement of α- and β-type globin genes from *B*. *pectinirostris*, we characterize the isoHb composition of adult red cells, and we integrate comparative genomic and phylogenetic analyses to shed light on the evolutionary origins of isoHb multiplicity in mudskippers.

## RESULTS

### Genomic organization of the globin gene clusters

We used a genome assembly of the blue-spotted mudskipper, *Boleophthalmus pectinirostris*, to characterize the physical organization of the globin gene clusters and we integrated analyses of conserved synteny with phylogenetic reconstructions to unravel the duplicative history of the mudskipper α- and β-type globin genes. We identified a total of four α-type globin genes and three β-type globin genes in the genome of *B. pectinirostris*. As in other teleosts examined to date, these genes were distributed between two unlinked clusters, designated ‘LA’ and ‘MN’ (the abbreviations refer to flanking genes that demarcate the 5’ and 3’ boundaries of each set of tandemly linked globin genes)(Fig. 1). The LA and MN gene clusters represent the paralogous products of the teleost-specific whole-genome duplication (TGD)(Opazo et al., 2013) that occurred ~350 million years ago (Meyer and Van de Peer, 2005). The LA cluster of *B. pectinirostris* contains two α-type globin genes (*Hba1-LA* and *Hba2-LA*, from 5’ to 3’) with two β-type globin genes interleaved between them in the opposite orientation (*Hbb2-LA* and *Hbb1-LA*), whereas the MN cluster contains three α-type globin genes (*Hba1-MN*, *Hba2-MN*, and *Hba3-MN*)(Fig. 2).

**Figure 1.**
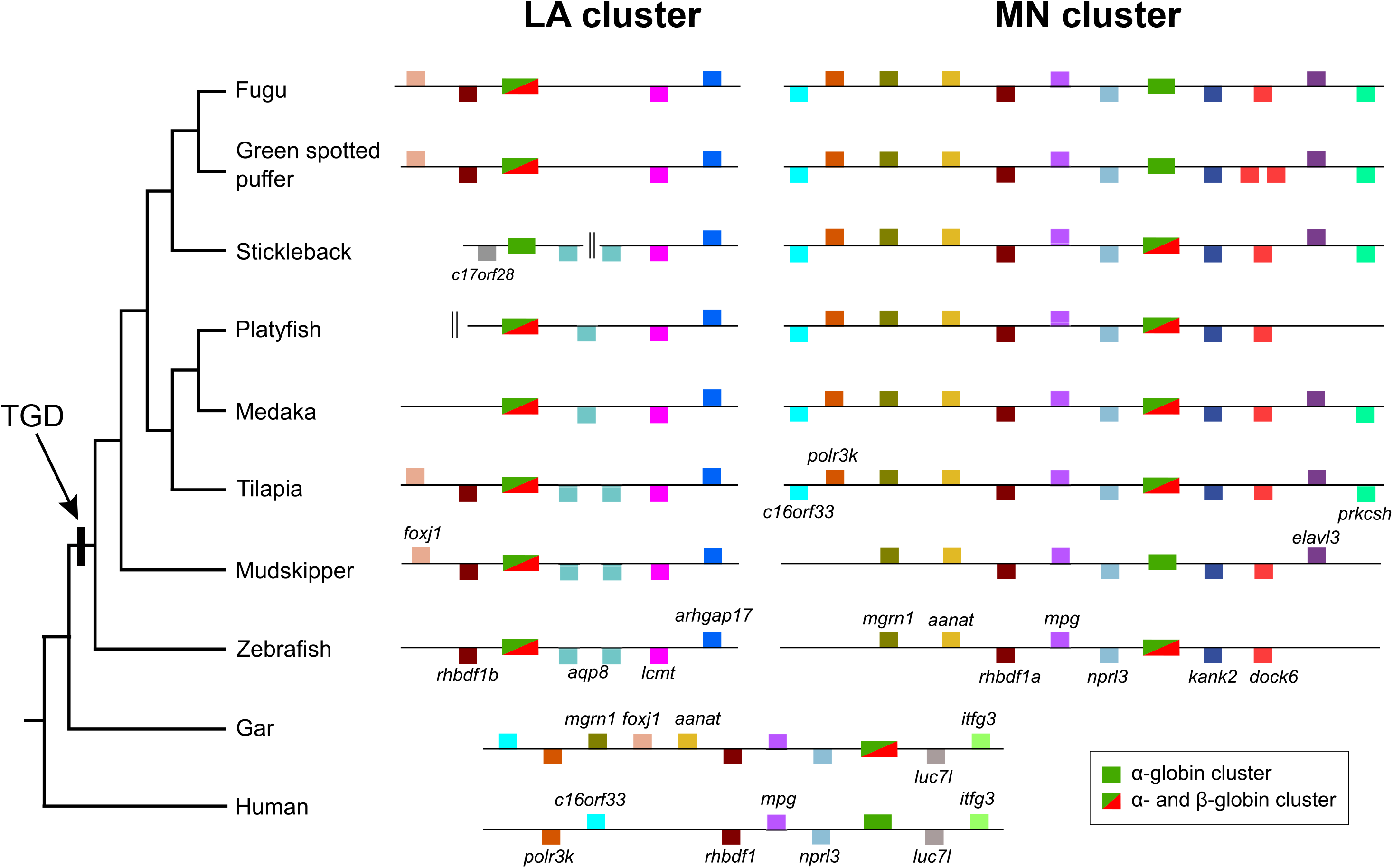
Unscaled representation of the genomic organization of the LA and MN globin gene clusters in the blue-spotted mudskipper, *Boleophthalmus pectinirostris*, and other representative ray-finned fish. The α-globin gene cluster of human is shown for comparison. As discussed in the text, the LA and MN gene clusters represent paralogous chromosomal segments that are products of the teleost-specific genome duplication (Opazo et al., 2013). Genes in the sense orientation are shown above the chromosome, those in the anti-sense orientation are shown below, and the globin gene clusters (which include genes in both orientations) are vertically aligned along the mid-line. The globin gene clusters of each species are shown in the same orientation as those in the zebrafish genome.

**Figure 2.**
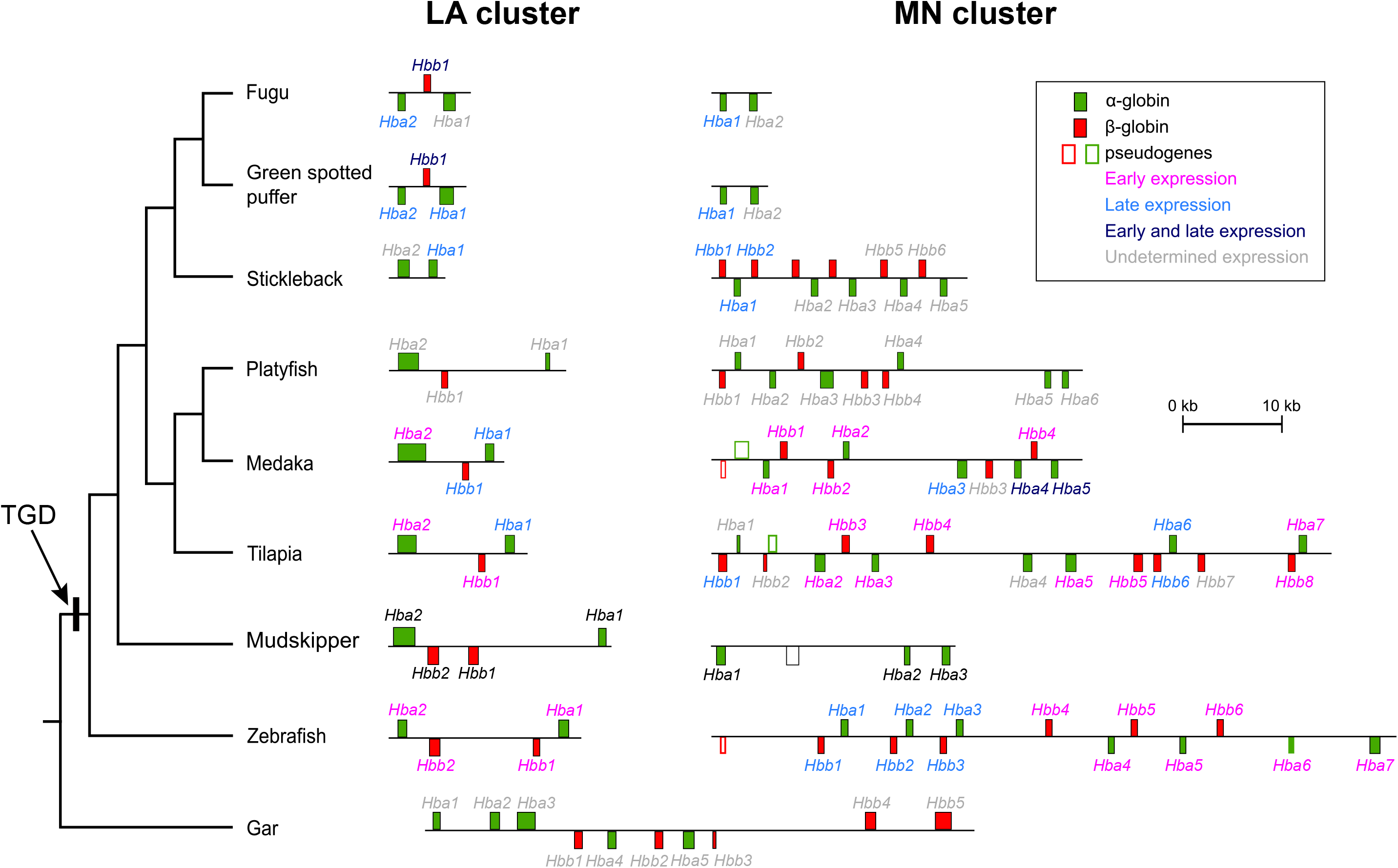
Genomic organization of the MN and LA globin gene clusters of the blue-spotted mudskipper, *Boleophthalmus pectinirostris*, and a representative set of other ray-finned fishes. As discussed in the text, the LA and MN gene clusters represent paralogous chromosomal segments that are products of the teleost-specific genome duplication (TGD), which occurred after the ancestral line of extant teleosts diverged from the ancestor of gars. The abbreviations LA and MN refer to flanking genes that delineate the 5’ and 3’ boundaries of each set of tandemly linked globin genes (Opazo et al., 2013). Globin genes in the sense orientation are shown above the chromosome and genes in the anti-sense orientation are shown below.

### IsoHb composition of *B. pectinirostris* red cells

Red cell lysates from *B. pectinirostris* were subjected to anion-exchange fast protein chromatography (FPLC), which revealed two major adult isoHbs, HbI and HbII, in approximately equal quantities and with similar anodic mobilities on native polyacrylamide gels. Tandem mass spectrometry (MS/MS) analysis of the FPLC-purified isoHbs revealed that each of them incorporate the same β-chain (product of the *Hbb1-LA* gene) but different α-chains, as HbI incorporates the product of *Hba1-LA* and HbII incorporates the product of *Hba1-MN* (Fig. 3A). The two α-globin genes, *Hba1-LA* and *Hba1-MN*, are members of the two separate TGD-derived gene clusters (Fig. 2). Given the antiquity of the TGD, it is not surprising that the two α-globin paralogs are highly divergent at the amino acid sequence level, differing at 54 of 141 residue positions (Fig. 3B).

**Figure 3.**
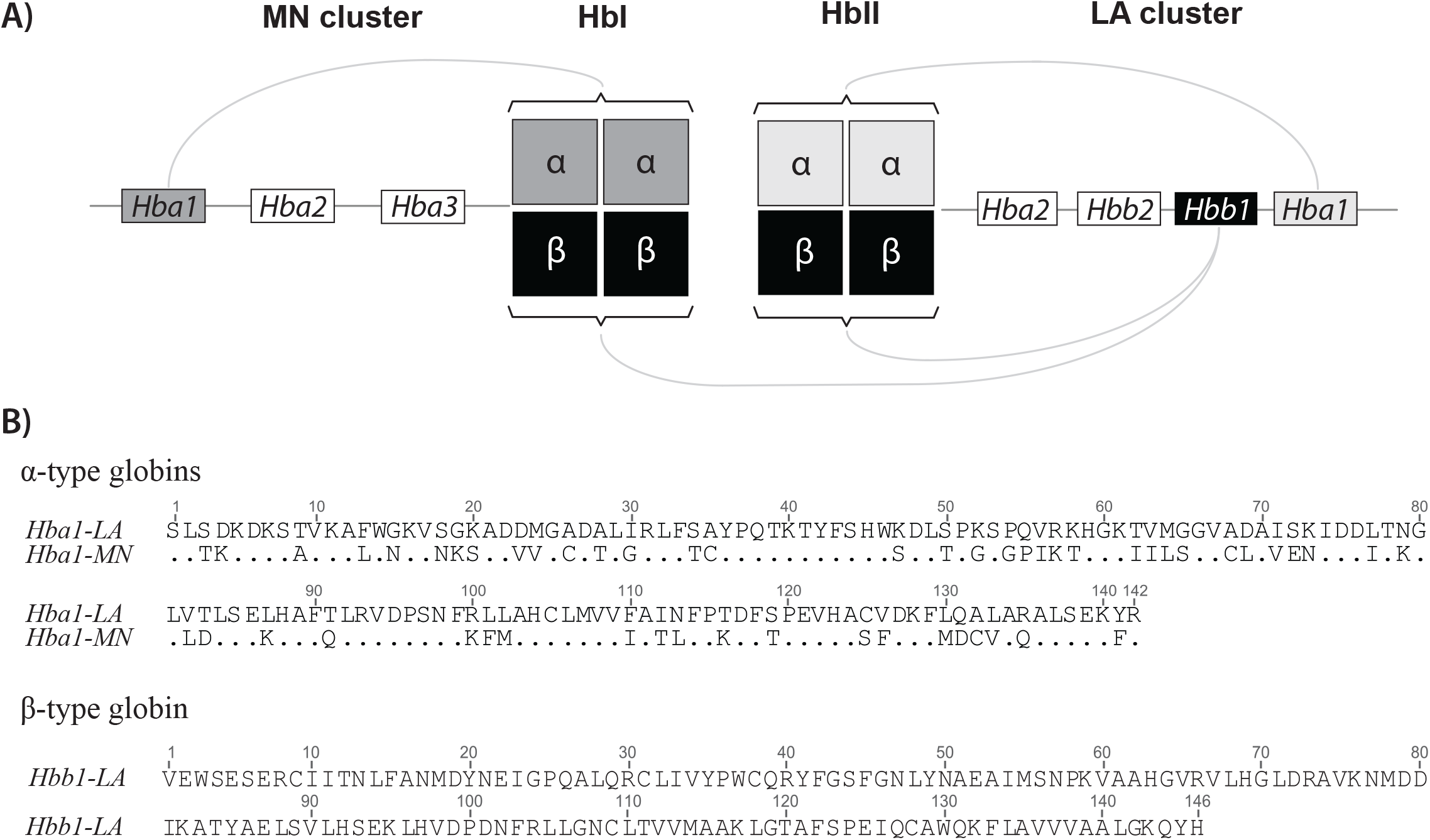
IsoHb composition of mudskipper isoHbs. A) Subunit composition of the two major isoHbs of the blue-spotted mudskipper, *Boleophthalmus pectinirostris*, as revealed by tandem mass-spectrometry. B) Amino acid sequences of adult α- and β-chain subunits.

### Phylogeny of α- and β-type globin genes

Consistent with the comparative genomic analysis of conserved synteny, phylogenetic analysis revealed that α- and β-type globin genes from a taxonomically diverse set of teleost species clustered into discrete LA and MN clades (Fig. 4). Within each main clade of LA- and MN-associated globin genes, the adult-expressed mudskipper globin genes were in some cases nested within subclades of genes from other species that are annotated as early-expressed embryonic globins. For example, within ‘LA Hba clade 2’ in the phylogeny of α-type globins (Fig. 4A), the adult-expressed *Hba1-LA* gene of mudskipper forms a subclade with the adult-expressed *Hba1-LA* gene from medaka (*Oryzias latipes*) and the early-expressed *Hba2-LA* gene from Nile tilapia (*Oreochromis niloticus*). Likewise, within the ‘LA Hbb clade 1’ in the phylogeny of β-type globins (Fig. 4B), the adult-expressed *Hbb1-LA* of mudskipper forms a subclade with the adult-expressed *Hbb1-LA* of medaka and the early-expressed *Hbb1-LA* of Nile tilapia, as well as the early- and late-expressed *Hbb1-LA* genes of green-spotted puffer (*Dichotomyctere nigroviridis*) and fugu (*Takifugu rubripes*).

**Figure 4.**
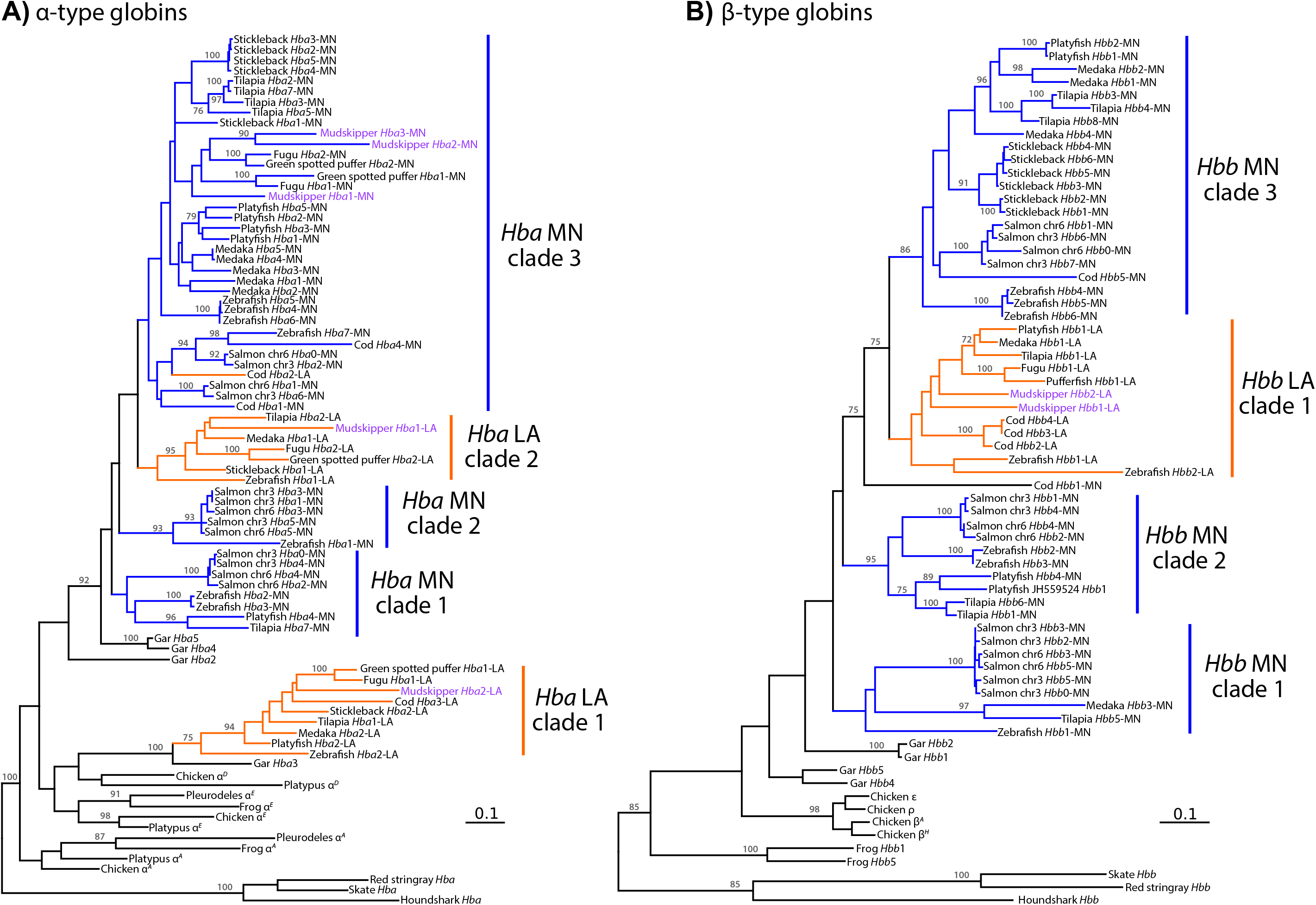
Maximum likelihood phylogenetic reconstruction depicting relationships among globin sequences of the blue-spotted mudskipper, *Boleophthalmus pectinirostris*, and a representative set of other ray-finned fish, cartilaginous fish, and tetrapods. Reconstructions were based on the coding sequences of α- and β-type globin genes (panels A and B, respectively). Only nodes with bootstrap support values greater than 70 are labeled. Branches are color-coded according to the location of the associated genes: MN-linked genes are shown in blue and LA-linked genes are shown in orange. The names of the mudskipper genes are written in purple. Scale bars for branch lengths denote the estimated number of nucleotide substitutions per site.

### O_2_-binding properties of mudskipper Hbs

Under identical experimental conditions, O_2_-equilibrium curves revealed that the two isoHbs exhibited virtually identical O_2_-affinities (as measured by *P*_50_, the *P*O_2_ at which Hb is 50% saturated) and cooperativities (as measured by *n*_50_, Hill’s cooperativity coefficient at half-saturation)(Fig. 5, Table 1). O_2_-affinities of the two isoHbs were reduced (i.e., *P*_50_’s were increased) in the presence of ATP but not in the presence of Cl^-^ ions alone (added as 0.1 M KCl)(Fig. 5, Table 1). Curves for both isoHbs measured in the joint presence of KCl and ATP revealed a slight decrease in *P*_50_ compared to curves measured with ATP alone, indicating that the binding of monovalent Cl^-^ ions interferes with the allosteric binding of ATP in the central cavity of T-state deoxyHb. The O_2_-affinities of both isoHbs exhibited pronounced sensitivities to changes in pH: Bohr factors (Δlog*P*_50_/ΔpH) were −0.79 and −0.81 for HbI and HbII, respectively.

**Figure 5.**
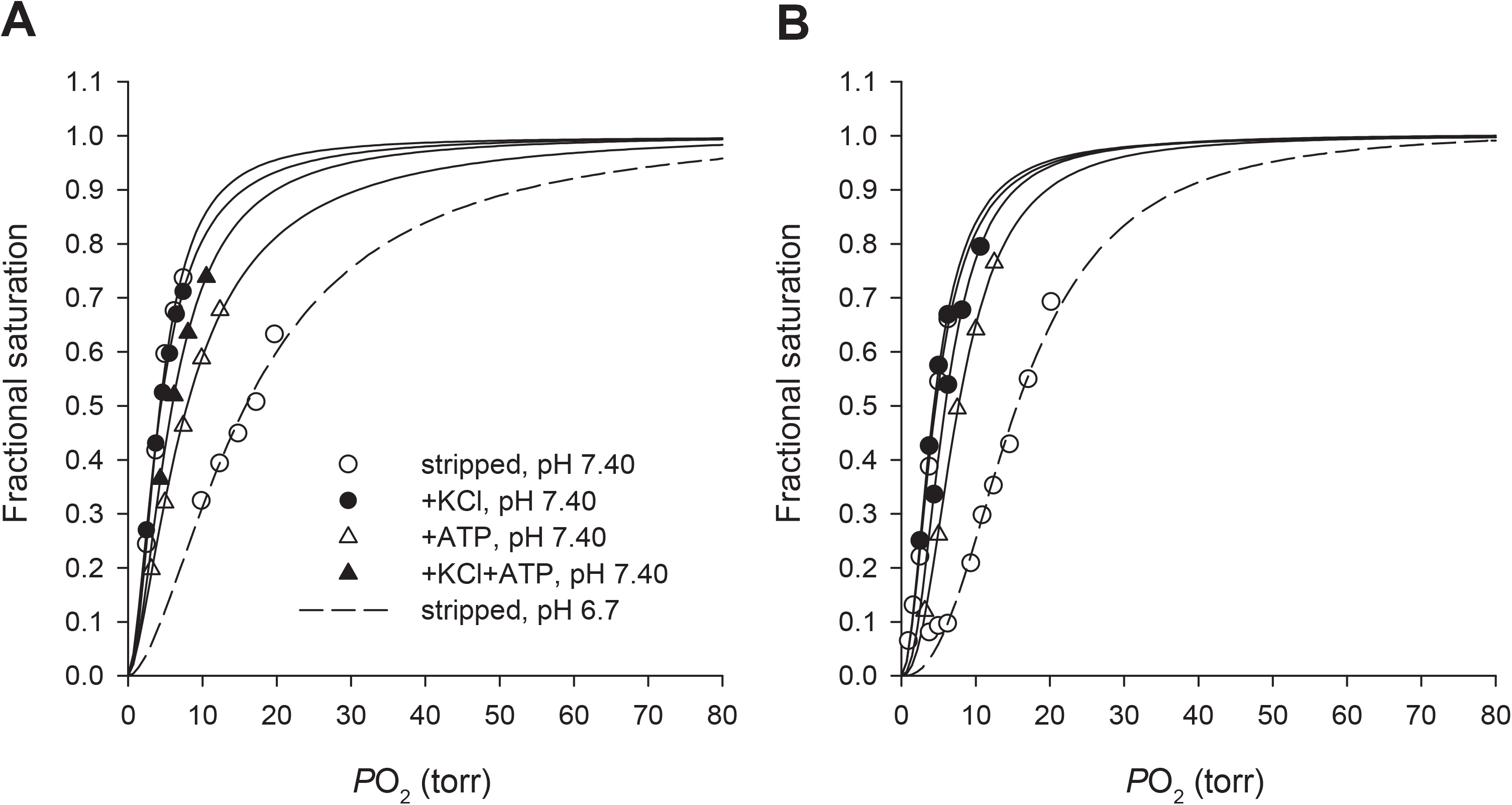
O_2_-equilibrium curves of purified mudskipper isoHbs. Curves for HbI (A) and HbII (B) were measured at 25°C in 0.1 M Hepes buffer in the absence (stripped) and presence of allosteric cofactors (Cl- ion [added as KCl]) and ATP (see *Materials and Methods* for details). To measure the Bohr effect, curves were measured under stripped conditions at pH 7.40 (solid lines) and at pH 6.67 (HbI) or 6.74 (HbII)(dashed lines).

**Table 1.**
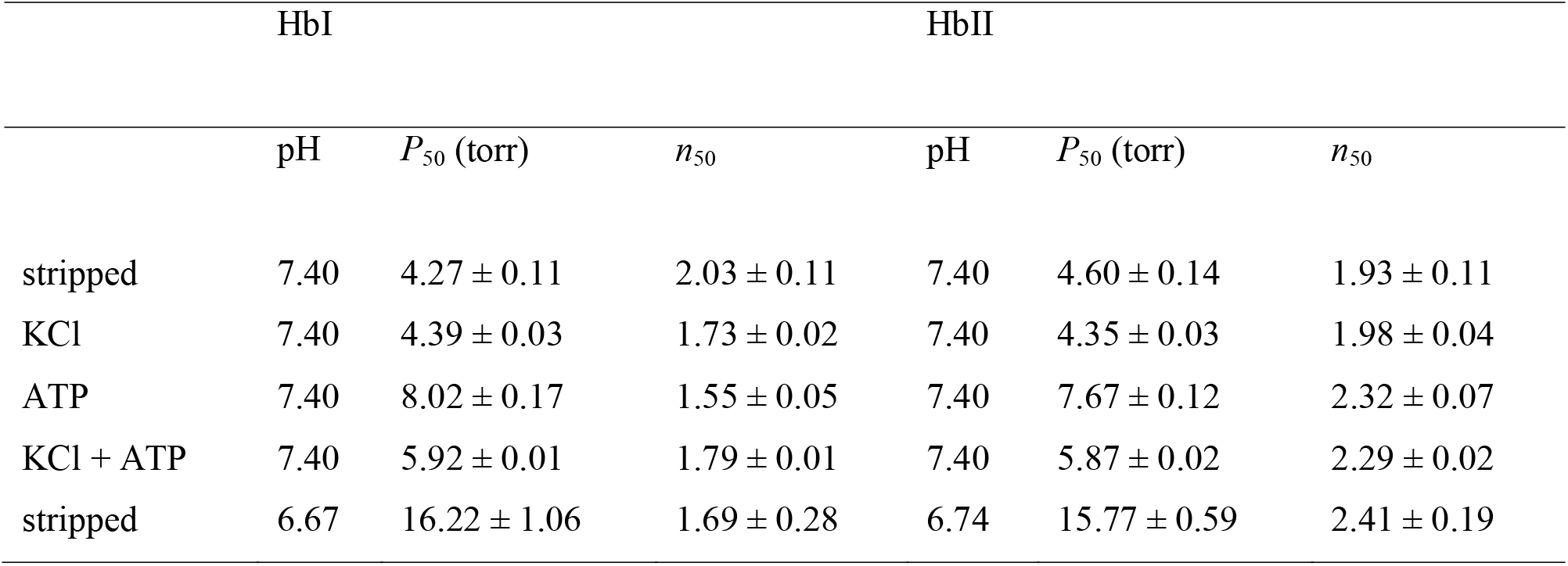
O_2_-binding properties of the two major isoHbs of the blue-spotted mudskipper, *Boleophthalmus pectinirostris* (means ± sem). *P*_50_ and *n*_50_ values were estimated from O_2_-equilibrium curves fitted by 4-9 saturation points (*r*^2^ =0.992-0.999).

|  | HbI |  |  | HbII |  |  |
| --- | --- | --- | --- | --- | --- | --- |
| | pH | $P_{50}$ (torr) | $n_{50}$ | pH | $P_{50}$ (torr) | $n_{50}$ |
| stripped | 7.40 | $4.27 \pm 0.11$ | $2.03 \pm 0.11$ | 7.40 | $4.60 \pm 0.14$ | $1.93 \pm 0.11$ |
| KCl | 7.40 | $4.39 \pm 0.03$ | $1.73 \pm 0.02$ | 7.40 | $4.35 \pm 0.03$ | $1.98 \pm 0.04$ |
| ATP | 7.40 | $8.02 \pm 0.17$ | $1.55 \pm 0.05$ | 7.40 | $7.67 \pm 0.12$ | $2.32 \pm 0.07$ |
| KCl + ATP | 7.40 | $5.92 \pm 0.01$ | $1.79 \pm 0.01$ | 7.40 | $5.87 \pm 0.02$ | $2.29 \pm 0.02$ |
| stripped | 6.67 | $16.22 \pm 1.06$ | $1.69 \pm 0.28$ | 6.74 | $15.77 \pm 0.59$ | $2.41 \pm 0.19$ |

## DISCUSSION

### Oxygenation properties of mudskipper Hbs

*B. pectinirostris* expresses multiple anodic isoHbs, each with a relatively high intrinsic O_2_-affinity, moderate sensitivity to organic phosphates, and a large Bohr effect – a “class I” profile, according to the classification scheme of Weber (1990). This same basic pattern of isoHb multiplicity has been well-documented in other teleosts, including facultative air-breathers and obligate water-breathers alike. The Hb system of *B. pectinirostris* appears functionally similar to that of the swamp eel (*Monopterus albus*), a facultative air-breather in which gas exchange is largely restricted to the buccopharyngeal cavity (Damsgaard et al., 2014; Damsgaard et al., 2015). Similar to *B. pectinirostris*, *M. albus* expresses multiple anodic isoHbs and no cathodic components. The two major isoHbs of *M. albus* exhibit high intrinsic O_2_-affinities (*P*_50(stripped)_ = 4.8-5.2 torr [25 °C, pH 7.7]), similar to values measured for the two isoHbs of *B. pectinirostris* (Table 1), but the mudskipper isoHbs exhibited a higher sensitivity to ATP and estimated Bohr coefficients were 2- to 3-fold larger when measured under identical buffer conditions. The mudskipper isoHbs have *P*_50_ values similar to those reported for the anodic isoHbs of hypoxia-tolerant, water-breathing teleosts such as carp (*Cyprinus carpio*)(Gillen and Riggs, 1972; Weber and Lykkeboe, 1978; Jensen et al., 2017) and European eel (*Anguilla anguilla*)(Fago et al., 1997). In contrast to the mudskippers, another facultative air-breathing fish, the Amazonian armored catfish (*Hoplosternum littorale*), exhibits a far more pronounced level of isoHb differentiation, with anodic and cathodic isoHbs that differ substantially in intrinsic O_2_-affinity, phosphate sensitivity, and Bohr effect (Weber et al., 2000).

The high Hb-O_2_ affinity of the mudskipper should promote branchial O_2_-uptake under conditions of aquatic hypoxia and – during terrestrial excursions – should promote transepithelial O_2_-uptake in the pharyngeal air-breathing structures. The potential drawback of an increased Hb-O_2_ affinity is that it hinders O_2_ unloading in the tissue capillaries, but this can be mitigated to some extent by the Bohr effect. The large Bohr effect of *B. pectinirostris* and some other facultative air-breathing fish may enhance O_2_ delivery to tissues during metabolic or respiratory acidosis upon switching from aquatic to aerial respiration, where O_2_ uptake may be less impaired. Likewise, liberation of Bohr protons upon Hb oxygenation would greatly facilitate CO_2_ excretion. Measurements on whole blood have revealed numerous facultative air-breathing fish species that exhibit pronounced Bohr effects, but there is extensive interspecific variation (Shartau and Brauner, 2014; Bayley et al., 2019; Mendez-Sanchez and Burggren, 2019). Given the especially large Bohr effects of the mudskipper isoHbs, it is worth noting that the β-chain C-termini (HC3) have His (Fig. 3B), which is known to play a major role in deoxygenation-linked proton-binding, whereas the cathodic isoHbs of other teleosts like *Hoplosternum* and *Anguilla* (which exhibit little or no Bohr effect, or even a reverse Bohr effect in the absence of organic phosphates) have non-ionizable Phe at the same residue position (Fago et al., 1995; Weber et al., 2000).

In addition to the hypothesis that obligate and facultative air-breathing fishes have evolved reduced Hb-O_2_ affinities compared to obligate water-breathers (Johansen and Lenfant, 1972; Johansen et al., 1978; Powers et al., 1979), it has been suggested that facultative air-breathing fishes may have evolved Hbs with reduced Bohr effects to cope with CO_2_ retention and/or exposure to hypercarbic water (Carter, 1931). Available data do not appear to support either hypothesis. Among ray-finned fish, lobe-finned fish, and amphibians, facultative air- and water-breathers tend to have higher blood-O_2_ affinities than obligate air-breathers, and facultative air-breathers tend to have slightly larger Bohr effects than members of the other two groups (Bayley et al., 2019). The high intrinsic O_2_-affinities and large Bohr effects that we measured for the purified mudskipper isoHbs appear to be consistent with both trends.

### Genomic insights into the origins of Hb multiplicity in mudskippers

Given that the two adult-expressed isoHbs of *B. pectinirostris* are present at approximately equimolar concentrations in the red cell, it seemed reasonable to expect that the two distinct α-chain subunits would be encoded by a tandemly linked pair of genes under the transcription control of the same *cis*-acting regulatory elements. Such a linkage arrangement could also help explain why the two isoHbs exhibit such similar O_2_-binding properties, since tandem duplicates often have similar coding sequences due to interparalog gene conversion (Hoffmann et al., 2008a; Hoffmann et al., 2008b; Runck et al., 2009; Storz et al., 2010; Gaudry et al., 2014; Natarajan et al., 2015; Signore et al., 2019). Contrary to this seemingly reasonable expectation, the analysis of conserved synteny revealed that the genes that encode the α-type subunits of HbI and HbII are members of unlinked, paralogous gene clusters that represent the duplicated products of the teleost-specific genome duplication (Figs. 1 and 2).

In the phylogenies of α- and β-type globins, the fact that adult-expressed mudskipper genes clustered with some embryonically expressed globin genes of other teleosts is not too surprising, as evolutionary changes in stage-specific expression during ontogeny have been well-documented in teleosts (Opazo et al., 2013) and in many tetrapods as well (Opazo et al., 2008; Hoffmann et al., 2010; Storz, 2016; Hoffmann et al., 2018; Storz, 2019).

### IsoHb multiplicity and lack of functional differentiation

In the case of mudskippers, there is no appreciable difference in the functional properties of co-expressed isoHbs in spite of extensive amino acid divergence between the alternative α-chain subunit isoforms (Fig. 3B). It therefore appears that the ability to switch between aquatic and aerial respiration does not necessarily require a division of labor between functionally distinct isoHbs that are specialized for O_2_ transport under different conditions. The adult-expressed isoHbs of many obligate water-breathing fish are known to exhibit much higher levels of functional differentiation than those of mudskippers, including quantitative differences in O_2_-affinity and qualitative differences in the mode of allosteric regulation, especially in relation to the magnitude of the Bohr effect and sensitivity to organic phosphates (Weber, 1990; Weber, 2000). In many teleost species, the presence of specific isoHbs with functional specializations such as the Root effect (an extreme form of pH-sensitivity that plays a key role in oxygen secretion and general tissue O_2_ delivery) exerts an important influence on physiological capacities (Berenbrink, 2007; Rummer et al., 2013; Randall et al., 2014). However, the absence of functional isoHb differentiation in amphibious mudskippers and other facultative air-breathers, and the often pronounced functional heterogeneity in the isoHbs of obligate water-breathers, supports the conclusions of several authors (Fyhn et al., 1979; Ingermann, 1997; Wells, 2009; Storz, 2019) that the overall diversity of co-expressed isoHbs in fish red cells is not generally a strong determinant of physiological versatility or ecological niche breadth.

## MATERIALS AND METHODS

### Sequence data and phylogenetic analyses

We annotated the full complement of α- and β-type globin genes in genome assembly of the blue-spotted mudskipper, *Boleophthalmus pectinirostris* (GenBank accession number: JACK00000000.1). For analyses of conserved synteny and phylogenetic relationships, we used genomic sequence data from a diverse set of bony fish and cartilaginous fish, including globin gene sequences that we annotated previously (Opazo et al., 2013; Opazo et al., 2015).

Protein coding sequences were translated and aligned using L-INS-i strategies in Mafft v7.3 (Katoh and Standley, 2013) and were then reverse-translated so that coding sequences were codon-aligned. We annotated several genes with truncated coding sequences (*Hbb3-MN* in platyfish [*Xiphophorus maculatus*], *Hbb2-MN* and *Hbb6-MN* in Nile tilapia [*Oreochromis niloticus*], and *Hba1* in spotted gar [*Lepisosteus oculatus*]),but we did not include such sequences in the phylogenetic analyses. We reconstructed separate phylogenies for the α- and β-globin genes using maximum likelihood (ML). After partitioning the alignment into codon positions, we performed the ML analyses in RAxML v 8.1.3 (Stamatakis, 2006) using the GTRGAMMA substitution model. Support for each node in the best tree was based on 1,000 bootstrap replicates.

### Blood sampling and characterization of isoHb composition

Blood was drawn from two individual specimens of *B. pectinirostris* in accordance with an animal handling protocol approved by the Animal Ethics Committee and the Institutional Review Board on Bioethics and Biosafety of BGI (certification number FT15029). Each sample was incubated for 5 min in a 5-fold volume of ice-cold buffer (10 mM Hepes, 0.5 mM EDTA, pH 7.75). Hemolysates were centrifuged to remove cell debris (30 min at 15000 g, 4 °C) and KCl was then added to the clear supernatant to a final concentration of 0.2 M (Jendroszek et al., 2018). The sample was then passed through a 5 ml PD-10 (GE Healthcare) desalting column equilibrated with 10 mM Hepes, 0.5 mM EDTA, pH 7.7. This desalting step removes endogenous organic phosphates, which are mainly ATP and GTP in fish red cells.

### Separation and purification of isoHbs

We purified individual isoHbs by means of anion-exchange fast-protein liquid chromatography (FPLC) using an Äkta Pure system (GE Healthcare). Stripped hemolysate was loaded on the prepacked HiTrap Q-HP column (GE Healthcare), equilibrated with 10 mM Hepes buffer (0.5 mM EDTA, pH 7.66) and separation of individual isoHbs was achieved using a linear gradient of 0-80% of 0.2 M NaCl, with a flow rate of 1 ml/min. Fractions were concentrated using Amicon Ultra centrifugal filters (10kDa) and were subsequently dialyzed against 10 mM Hepes, 0.5 mM EDTA, pH 7.7, at 4 °C, to remove NaCl. Purity of the isoHb fractions was verified by means of native polyacrylamide gel electrophoresis (PAGE) using a PhastSystem (GE Healthcare) in 10-15% gradient precast gels. Purified samples were stored at −80 °C at a 1 mM heme concentration.

### Identification of IsoHb subunit composition

The subunit composition of each eluted Hb fraction was determined by tandem mass spectrometry (MS/MS). For the MS/MS analysis, eluted fractions were separated on a 20% SDS-PAGE gel and were stained with Coomassie brilliant blue. The individual α- and β-chain subunits of the purified isoHbs were excised and digested with trypsin. Following previously described protocols (Revsbech et al., 2013; Storz et al., 2015), peptide mass fingerprints derived from MS/MS were queried with a custom database that included amino acid sequences from all embryonic and adult-expressed α- and β-type globin genes that we annotated from the *B. pectinirostris* genome assembly. The queries were conducted with the Mascot data search system (Matrix Science, version 1.9.0, London, UK). Search parameters included no restriction on protein molecular weight or isoelectric point, and methionine oxidation was allowed as a variable peptide modification. Mass accuracy settings were 0.15 Da for peptide mass and 0.12 Da for fragment ion masses. We identified all significant protein hits that matched more than one peptide with *P* < 0.05.

### Measurement of Hb-O_2_ equilibria

We measured O_2_-equilibrium curves of purified isoHbs at 25°C in 0.1 M Hepes buffer (pH 7.40) in the absence (stripped) and presence of 0.1 M KCl and ATP (0.75 mM), and in the simultaneous presence of KCl and ATP. Heme concentration for each Hb solution was 0.3 mM. To measure the Bohr effect, we took replicate measurements of stripped isoHbs under identical conditions at pH 6.7 and 7.4. The final pH of Hb solutions used in each experiment was measured at 25°C using an InLab Micro pH electrode (Mettler Toledo). We measured O_2_-equilibria of 3 µl thin-film samples in a modified diffusion chamber where absorption at 436 nm was monitored during stepwise changes in the equilibration of N_2_/O_2_ mixtures generated by precision Wősthoff gas-mixing pumps (Weber, 1992; Grispo et al., 2012; Weber et al., 2013; Natarajan et al., 2015; Natarajan et al., 2016). We estimated values of *P*_50_ and *n*_50_ (Hill’s cooperativity coefficient at half-saturation) by fitting the Hill equation *Y*=*P*O_2_^*n*^ /(*P*_50_^*n*^+*P*O_2_^*n*^) to the experimental O_2_ saturation data by means of nonlinear regression (*Y*=fractional O_2_ saturation; *n*, cooperativity coefficient). The nonlinear fitting was based on 5–8 equilibration steps between 30% and 70% oxygenation. Free Cl^-^ concentrations were measured with a model 926S Mark II chloride analyzer (Sherwood Scientific Ltd, Cambridge, UK).

## Author contributions

Conceptualization: J.F.S., A.F., F.G.H. B.V.; Experiments and data analysis: C.N., M.G.K., M.V.; Sample acquisition: X.Y., B.V.; Writing – original draft: J.F.S., A.F.; Writing – review & editing: all authors; Funding acquisition: J.F.S.

